# In vitro evolution reveals primordial RNA-protein interaction mediated by metal cations

**DOI:** 10.1101/2021.08.01.454623

**Authors:** Valerio G. Giacobelli, Kosuke Fujishima, Martin Lepšík, Vyacheslav Tretyachenko, Tereza Kadavá, Lucie Bednárová, Petr Novák, Klára Hlouchová

## Abstract

RNA-peptide/protein interactions have been of utmost importance to life since its earliest forms, reaching even before the last universal common ancestor (LUCA). However, the ancient molecular mechanisms behind this key biological interaction remain enigmatic because extant RNA-protein interactions rely heavily on positively charged and aromatic amino acids that were absent (or heavily under-represented) in the early pre-LUCA evolutionary period. Here, an RNA-binding variant of the ribosomal L11 C-terminal domain was selected from a ∼10^10^ library of partially randomized sequences, all composed of 10 prebiotically plausible canonical amino acids. The selected variant binds to the cognate RNA with a similar overall affinity although it is less structured in the unbound form than the wild-type protein domain. The variant complex association and dissociation are both slower than for the wild-type, implying different mechanistic processes involved. The profile of the wild-type and mutant complex stabilities along with MD simulations uncover qualitative differences in the interaction modes. In the absence of positively charged and aromatic residues, the mutant L11 domain uses bridging ion (K^**+**^/Mg^2**+**^) interactions between the RNA sugar-phosphate backbone and glutamic acid residues as an alternative source of stabilization. This study presents experimental support to provide a new perspective on how early protein-RNA interactions evolved, where the lack of aromatic/basic residues was compensated by acidic residues plus metal ions.

## Introduction

Protein-RNA complexes are ubiquitous in modern life and are essential to many stages of the cell cycle and metabolism. The abundance of ribonucleoprotein (RNP) complexes generally increases with organism complexity (Balcerak et al. 2019). However, it is thought that peptide-RNA interactions date to life’s origins and have changed substantially over time (Hsiao et al. 2009; Lupas and Alva 2017; Vázquez-Salazar and Lazcano 2018).

Today’s proteins and RNAs can act as independent functional and structural entities and their interaction is often transient. On the other hand, early oligopeptides and RNA are thought to have served as mutual cofactors and scaffolds, necessitating strong interactions (Lupas and Alva 2017; Vázquez-Salazar and Lazcano 2018). Structural analyses of today’s natural and artificially evolved protein-RNA interactions show that proteins build these interactions by exploiting positively charged and aromatic amino acids (Blanco et al. 2018; Srivastava et al. 2018). However, these amino acids (Lys, Arg, His, Phe, Tyr, Trp) were not likely available or extremely scarce in the prebiotic environment (Trifonov 2000; Higgs and Pudritz 2009; Granold et al. 2018). As a result, the specific nature of the interaction between early peptides and RNA remains an open question in the field. Currently, there are two major hypotheses concerning the nature of early peptide-RNA binding. The first scenario holds that cationic amino acids were indispensable and their biosynthesis preceded the evolution of protein-RNA interaction (Blanco et al. 2018). Some studies suggest that the lack of canonical cationic amino acids in the early environment could be compensated by the presence of more abundant positively charged non-canonical amino acids (including some with positive side chains) originating from various prebiotic sources of organic material (Zaia et al. 2008; Cleaves 2010; Burton et al. 2012). Give that oligopeptides likely formed preceding the emergence of ribosomal translation, it follows that their composition would have reflected the composition of the prebiotic broth including non-canonical cationic amino acids, such as 2,4-diaminobutyric acid or ornithine (Raggi et al. 2016; Vázquez-Salazar and Lazcano 2018; Longo et al. 2020). An alternative scenario of early peptide-RNA interaction proposes that the electrostatic interaction of RNA with peptides relied on anionic (Glu and Asp, both prebiotically abundant) instead of cationic amino acids. In this model, the interaction would be mediated *via* metal ions (Raggi et al. 2016; Vázquez-Salazar and Lazcano 2018). The two hypotheses are not mutually exclusive and may also represent different evolutionary stages or combinations of different interaction types over the evolutionary timeline. At the same time, there is a lack of experimental support for both of them. To test whether strong protein-RNA interactions are conceivable in the absence of positively charged and aromatic amino acids, we performed a high-throughput variant analysis of a highly conserved RNA-binding domain of the ribosomal protein L11. We designed combinatorial libraries that comprised of (i) the prebiotically plausible subset of 10 and (ii) 14 “mid-stage” canonical amino acids, and selected RNA-binding mutants using mRNA display. Detailed characterization of viable variants from each of the two libraries unambiguously support that strong and specific protein-RNA interaction is possible in the absence of cationic and aromatic amino acids with the aid of metal ions. This mechanism of interaction confirms the feasibility of early RNA-protein interaction and brings attention to alternative RNA-protein engineering strategies.

## Results

### Design of L11 RNA binding domain combinatorial mutant libraries

To test the capacity of RNA-protein interaction in the absence of cationic and aromatic amino acids, we selected a highly conserved complex of the C-terminal RNA binding domain of *Bacillus stearothermophilus* ribosomal protein L11 (CL11) and a 58-nt domain of the large subunit ribosomal RNA (58rRNA) fragment as a selection target (Fig. 1) (Conn et al. 2002). Two variant libraries were designed following the ranking of prebiotic amino acid composition of early genetic code, as derived by the meta-analysis of Higgs and Pudritz (Higgs and Pudritz 2009). The “early” (E) library randomizes the native CL11 sequence by subsets of Gly, Ala, Asp, Glu, Val, Ser, Ile, Leu, Pro, Thr (Fig. 1A) while the “mid-stage” (M) library includes all the wild-type cationic and aromatic amino acids (Lys, Phe, Arg, His) as visualized in Fig. 1B. Sequences in the E library are therefore composed exclusively of the “early” amino acids with 26 % of the positions in the native sequence randomized. The substitution pools at each position in the were selected based on (i) environment-specific amino acid substitution tables that have been developed on the basis of acceptability of mutations within folded proteins by comparative analysis of homologous proteins, and (ii) multiple alignments of the protein family (Fig. 1C). This approach limited the size of the variant libraries E and M to ∼10^10^ and ∼10^6^, respectively. To represent all the desired substitution combinations in the E and M combinatorial libraries, degenerate codon DNA sequences for the library synthesis were designed using the SwiftLib tool (Jacobs et al. 2015).

**Figure 1.**
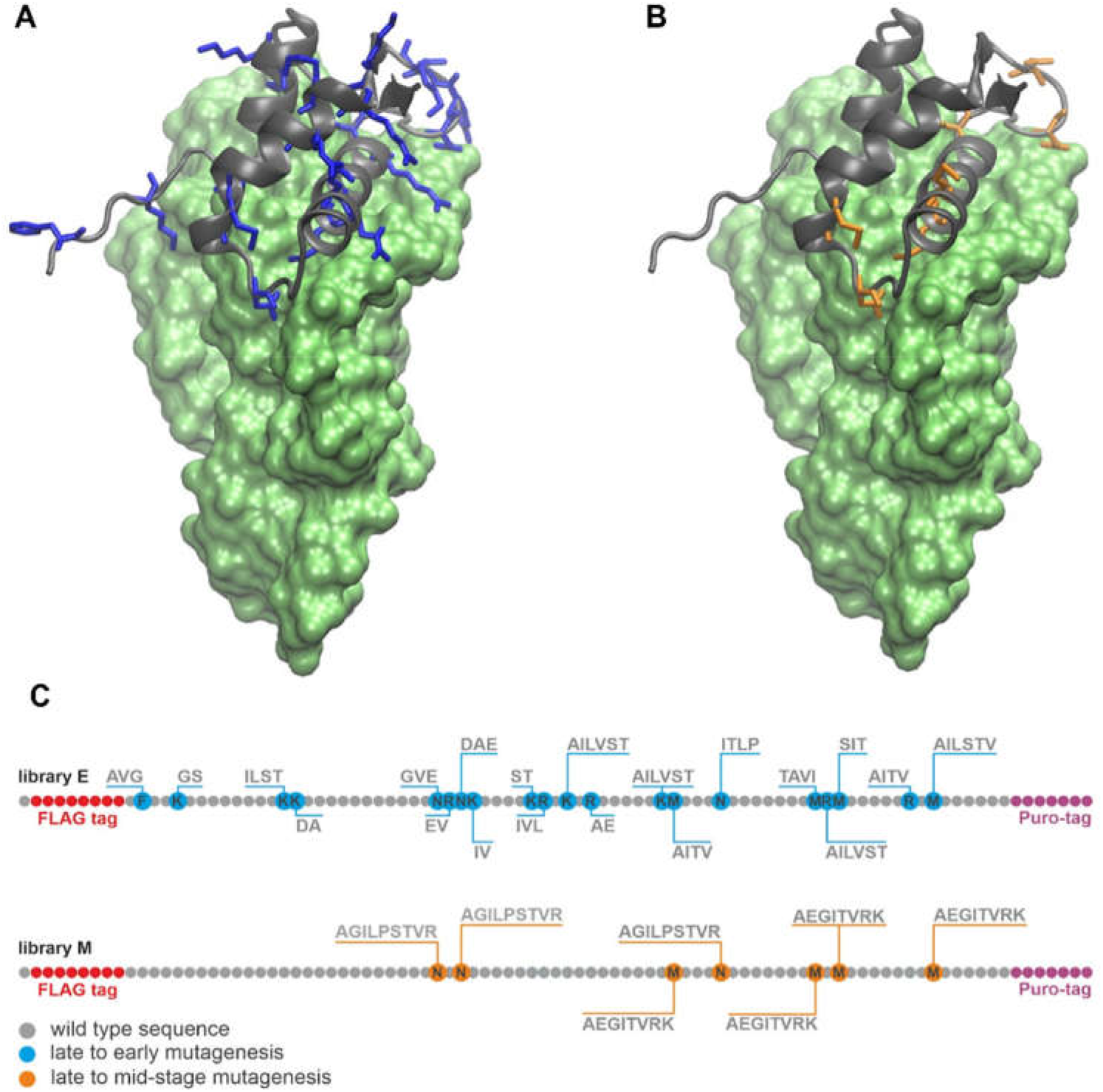
Design of CL11 variant libraries. Crystal structure of the CL11-58rRNA complex: (A) CL11: grey cartoon; CL11 substitution spots late to early amino acids: blue sticks; 58rRNA: green surface; (B) CL11: grey cartoon; CL11 substitution spots late to mid-stage amino acids: orange sticks; 58rRNA: green surface; (C) Randomization scheme for the CL11 -E and -M libraries.

### mRNA display of CL11 combinatorial libraries and RNA binding selection

The M and E oligonucleotide libraries were synthesized from two overlapping ssDNA oligonucleotides, including all the necessary sequence motifs for transcription and addition of the FLAG tag to the N’-terminal part of the protein product (see Fig. S1A). The oligonucleotide libraries were incorporated into a genotype-phenotype linked construct for mRNA display using a previously published protocol (Fig. S1B) (Reyes et al. 2021). Upon transcription and translation of the constructs, the best binding variants were selected against an immobilized 58rRNA, performing 10 and 16 rounds of the selections for libraries M and E, respectively. These numbers of mRNA display and selection rounds achieved distinct sequence consensus patterns at the randomized positions (Fig. 2). Interestingly, the E library reached an acidic pattern at two regions that were of basic character in the wild-type sequence, specifically “SD” in place of “KK”, and a “EVEV” motif in place of “NRNK” (Fig. 2A).

**Figure 2.**
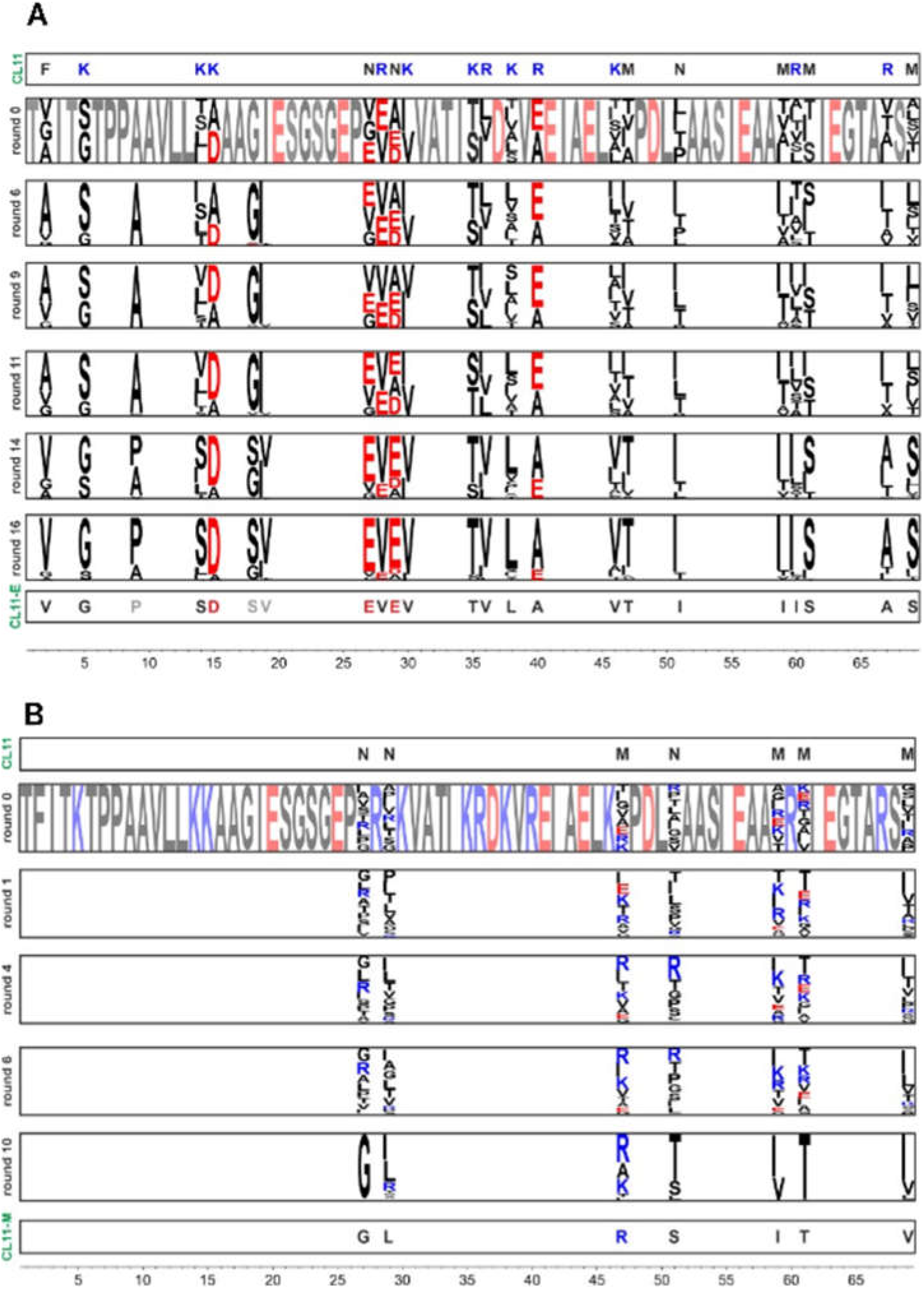
Characterization of CL11, CL11-M and CL11-E variants and their binding to 58rRNA. (A) CD spectra of CL11 (black), CL11-M (red) and CL11-E (green) in 30 mM Tris, 20 mM MgCl_2_ and 175 mM KCl at pH 7.9. (B) EMSA assay where equimolar concentration of f58rRNA target was incubated with the different protein variants. Free f58rRNA was used as a negative control. (C) Kinetic parameters of CL11, CL11-M and CL11-E binding to 58rRNA determined by SPR. (D) Protein-RNA complex stability at different temperatures, pHs and salts presence. Green: no protein is detected from western blot; increasing red intensity: increasing amount of protein detected on Western blot of the pull-down flow-through fraction (for original Western blots see Fig S4B)

The most enriched sequences from each library (CL11-E and CL11-M, representing 4.28 and 9.17 % of total read count respectively) were selected for detailed characterization (Fig. S2). The most abundant sequence from the E library (CL11-E) contained three additional mutations at fixed amino acid positions in the original library (Asp9Pro, Gly18Ser and Ile19Val). These mutations were likely introduced during the reverse transcription PCR step by the polymerases (Fig. 2A).

### CL11, CL11-M and CL11-E structural and binding characterization

Synthetic genes of the CL11 wild-type protein and CL11-E and CL11-M variants were subcloned with N’-terminal histidine tag and expressed in *Escherichia coli* OverExpress C41(DE3) pLysS cells. The purified protein samples were used to estimate the secondary structure content by circular dichroism (CD). The CL11 wild-type spectra are indicative of high α-helical structure content (∼45%) by the minima at ∼208 and ∼222 nm (Fig. 3A and Table S1). In contrast, both the CL11-E and CL11-M variants have significantly less secondary structure content. Both mutants seem to be mostly disordered as indicated by a profound negative peak at 202 nm in the CD spectrum and with an isosbestic point at 211 nm in temperature dependent CD spectra (Fig. S3A) (Uversky 2009).

**Figure 3.**
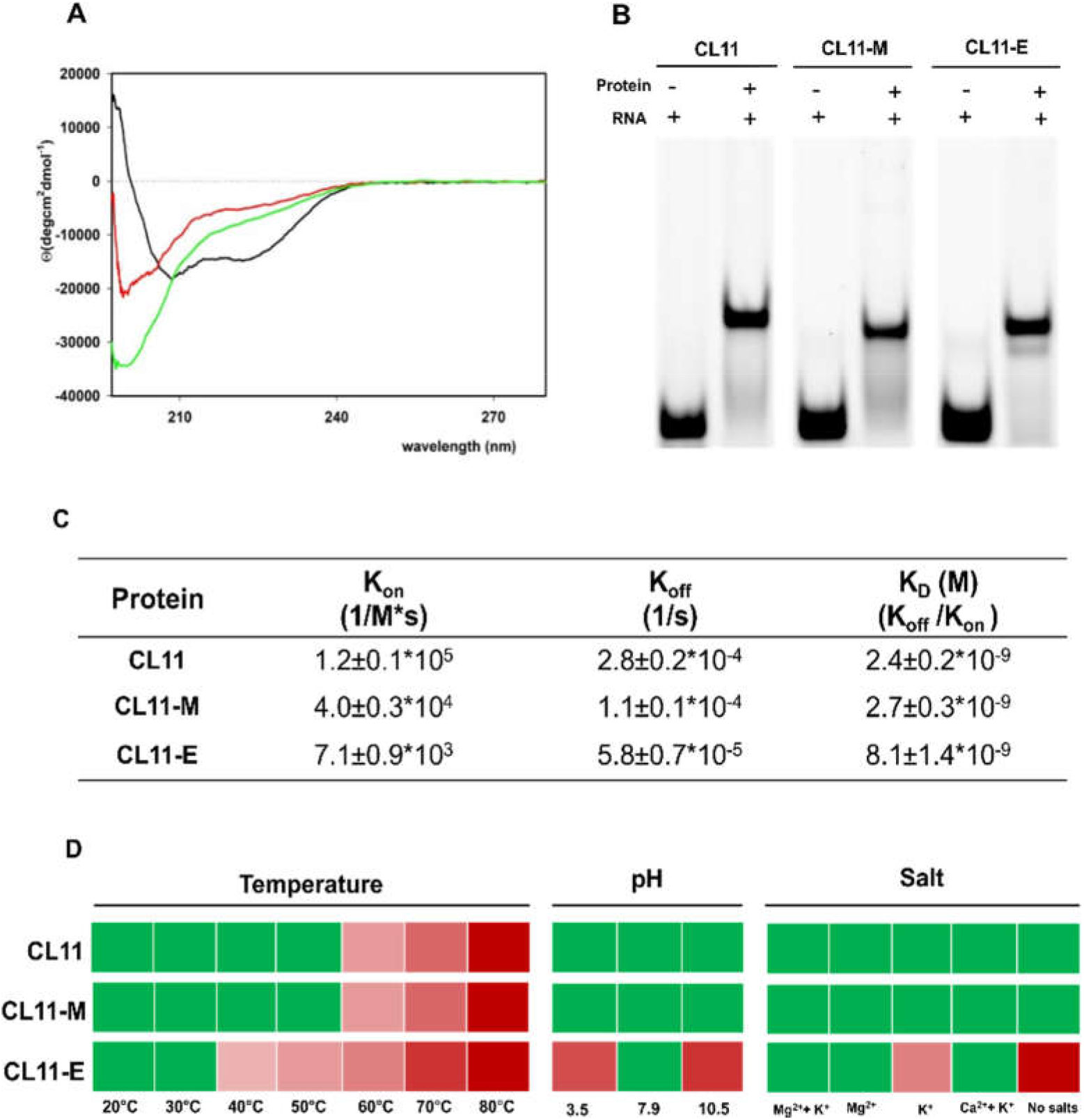
Characterization of CL11, CL11-M and CL11-E variants and their binding to 58rRNA. (A) CD spectra of CL11 (black), CL11-M (red) and CL11-E (green) in 30 mM Tris, 20 mM MgCl_2_ and 175 mM KCl at pH 7.9. (B) EMSA assay where equimolar concentration of f58rRNA target was incubated with the different protein variants. Free f58rRNA was used as a negative control. (C) Kinetic parameters of CL11, CL11-M and CL11-E binding to 58rRNA determined by SPR. (D) Protein-RNA complex stability at different temperatures, pHs and salts presence. Green: no protein is detected from western blot; increasing red intensity: increasing amount of protein detected on Western blot of the pull-down flow-through fraction (for original Western blots see Fig S4B).

The binding propensity of the purified protein to 58rRNA was confirmed using the electrophoretic mobility shift assay (EMSA). All the wild-type sequences (CL11), as well as the selected variants CL11-E and CL11-M, exhibited an electrophoretic shift on a native PAGE gel in the presence of f58rRNA (Fig. 3B). Dissociation constants of these binders were measured using surface plasmon resonance (SPR). The biotinylated 58rRNA was immobilized on the SPR chip by a neutravidin-biotin interaction and neutravidin alone was used as a negative control (Fig. S4A). While the *K*_*D*_ constants of both the CL11-E and CL11-M variants are comparable to that of the wild type CL11 protein (8.1×10-9 M and 2.7×10-9 M *vs*. 2.4×10-9 M, respectively), the *k*_*on*_ and *k*_*off*_ values indicate different kinetics of the RNA binding of these variants (Fig. 3C).

To test the stability of the protein-RNA complexes under different conditions, additional pull-down experiments were performed with the purified proteins against the immobilized 58rRNA, similar to the binding selection procedure during the mRNA-display protocol. The protein-RNA complexes were incubated at different temperatures, pH and salt contents (Fig. 4C). The flow-through samples, indicating the unstable protein fractions of the complexes, were analyzed by Western blot (Figs. 3D and S4B). While the complex stabilities are the same in all the tested conditions for CL11 and CL11-M, the CL11-E-RNA complex is destabilized at lower temperatures (40 to 50 ºC), extreme pH conditions and significantly affected by lack of Mg^2^**+** and K**+** ions. In addition, the CD spectra of the uncomplexed proteins were collected at the same pH and “no salts” condition to resolve the potential reasons of the complex instability (Fig. S3B). While the binding characteristics of the CL11-M mutant seem to be generally preserved with respect to the wild-type, the structural and RNA-binding properties of the CL11-E mutant prompted a further investigation to characterize the intriguing nature of this interaction.

**Figure 4.**
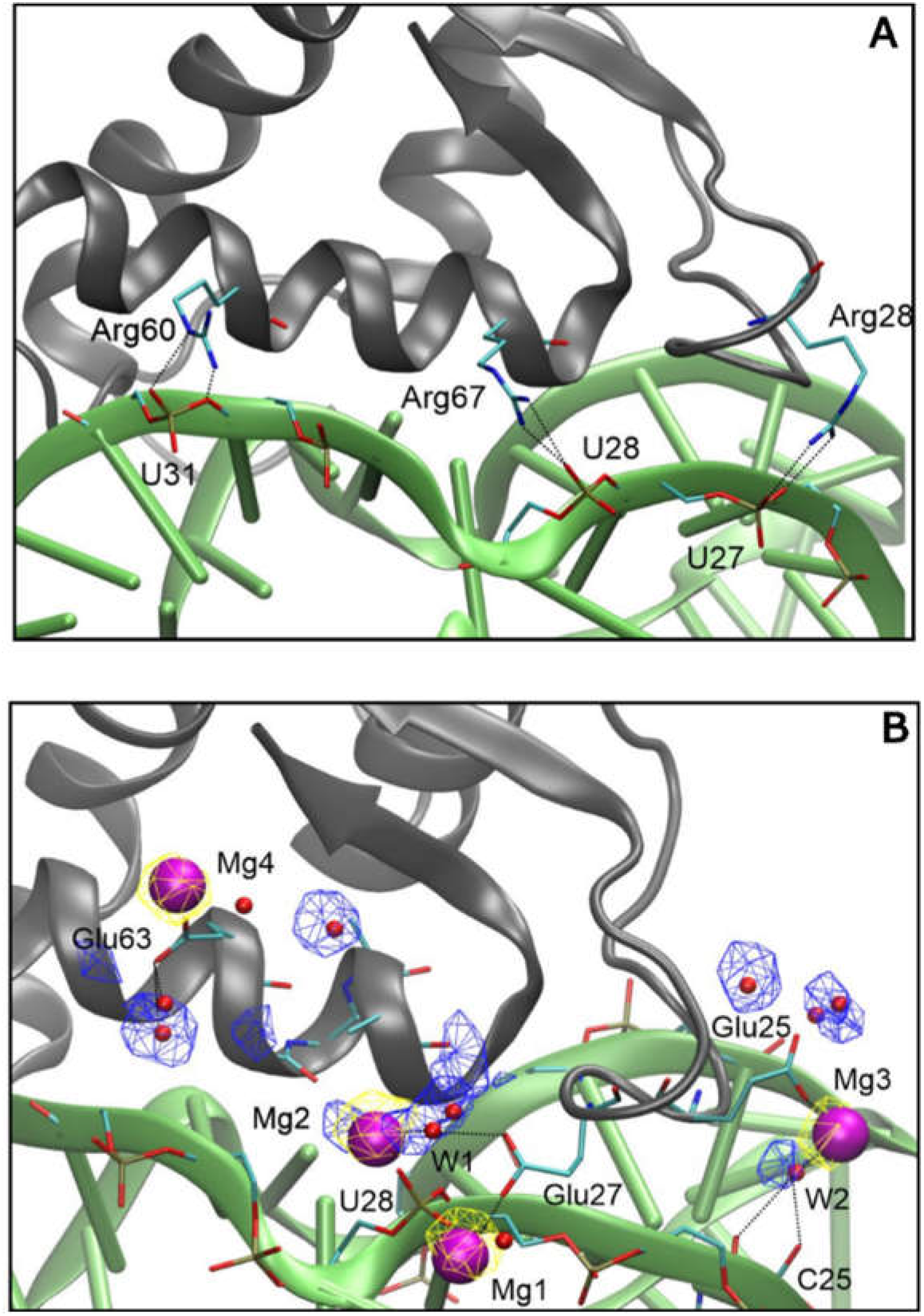
Molecular details of RNA-protein interaction. Snapshots from the last 500 ns of MD simulations showing the CL11/CL11-E proteins (grey cartoon) with important residues in sticks (cyan: C, blue: N, red: O, golden: P, hydrogens not shown), 58rRNA (green cartoon) with important residues in sticks, Mg^2+^ ions (purple spheres) and water molecules (red spheres). Yellow and blue meshes show the conserved sites for Mg^2+^ and water, respectively. Black dotted lines indicate H-bonding or metal coordination. (A) CL11 protein-RNA, (B) CL11-E protein-RNA.

We reasoned it would be difficult or impossible to crystallize the CL11-E-58rRNA complex; hence, an ion mobility experiment was performed to characterize overall structural properties. Both CL11 and CL11-E were incubated with 58rRNA and each complex was transferred to electrospray compatible buffer for an ion-mobility measurements. The CL11-58rRNA and CL11-E-58rRNA complexes revealed the 9^-^ion signal at m/z 3110 and 3030, respectively. The collision cross section was calculated 2130 Å^2^ and 1960, 2080, 2300 Å^2^ for ions at m/z 3110 and 3030 pointing out 1960 and 2080 Å^2^ conformers as a major fraction of the CL11-E-58rRNA sample. Since these CL11-E-58rRNA conformers differed from CL11-58rRNA complex by 50 and 170 Å^2^, we can conclude that both particles (CL11-58rRNA and CL-11-E-58rRNA complexes) kept similar shape in solution (Fig. S5).

### CL11 and CL11-E computational structural analysis

Using the previously published X-ray structure of the CL11-58rRNA complex (PDB code 1HC8), an initial model for CL11-E was prepared by generating the appropriate mutations in PyMol. The wild-type and the CL11-E apo-proteins, as well as their 58rRNA complexes, were prepared for molecular dynamics (MD) simulations to compare (i) the protein stabilities and (ii) the RNA-protein interface interactions. Throughout 2 µs MD, the behaviour of the proteins (in both complexed and uncomplexed forms) was relatively stable with average root-mean-square deviations (RMSD) of protein backbone atoms in secondary structure elements ranging from 0.6 to 1.1 Å. There were minor rearrangements around 1.3 µs and in the first 500 ns of the CL11-58rRNA and CL11-E-58rRNA complexes, respectively (Fig. S6). Nevertheless, the last 500 ns were stable in MD of all the systems studied and were thus used in further analyses. To investigate the nature of the CL11 *vs* CL11-E interaction with the 58rRNA, direct hydrogen bonds (H-bonds) were measured. 30 direct protein-RNA H-bonds (with an average occupancy of 63%) vs. 17 (with an average occupancy of 53%) were recorded for CL11 and CL11-E with the 58rRNA, respectively (Tab. S2); that is, CL11 makes nearly twice as many direct H-bonds with the RNA as compared to CL11-E. In 70% of cases, CL11 forms H-bonds via protein side chains (12 in total from Arg28, Arg60 and Arg67, Fig. 4A), while in the case of CL11-E, a slight majority (53%) of H-bonds are formed through backbone atoms. In addition, several water bridges were identified at the binding interface. In the case of CL11, these were mostly via Lys5, Lys14, Lys46, Arg 60 and Arg 67, while for CL11-E the side chains of Ser21, Ser61, Asp49 and the backbone of Ala10, Gly22 and Gly24 were the major residues involved. Further, metal (K^+^/Mg^2+^) ion sites were observed to bridge the protein and the RNA but only in the simulation of the CL11-E-58rRNA complex. Four K^+^ ions found their way to the areas of high electrostatic potential, *i*.*e*. close to carboxylates of protein Glu side chains and to the phosphates of RNA backbone. The stability of these sites in MD was increased upon exchanging K^+^ with Mg^2+^. Mg^2+^ ions were coordinated either directly by the RNA (U28 in Mg1 site, Fig. 4B) or the protein side chains (Glu25, Glu27 or Glu 63 in Mg3, Mg1 and Mg4 sites, respectively) or indirectly via water bridges (W1, W2 in Mg2 and Mg3 sites, respectively). It should be noted that Glu27 is a mutation with respect to the wild-type CL11 and that Mg4 does not provide an interaction with the RNA. Taken together, based on the MD simulation data, we propose that three Mg^2+^ mediate unique CL11-E specific interactions with 58rRNA.

## Discussion

Our experiments show that protein binders of RNA can be found in sequences composed of a significantly reduced set of canonical amino acid alphabet. Using mRNA display, we selected rRNA binders from variant libraries of the ribosomal L11 C-terminal domain that was composed of (i) the prebiotically plausible subset “E” of 10 canonical amino acids (lacking both positively charged and aromatic residues) as well as (ii) a reduced evolutionary mid-stage “M” alphabet of 14 amino acids (where all the positively charged and aromatic amino acids of the L11 C-terminal protein sequence were conserved).

Extant RNA-protein interaction relies substantially on the presence of positively charged and aromatic amino acids (Blanco et al. 2018). However, these amino acids were heavily underrepresented in the early stages of life’s origins, while negatively charged amino acids were quite abundant (Trifonov 2000; Higgs and Pudritz 2009). Hence, our finding of a robust RNA-protein interaction in the absence of aromatic/basic residues may represent an early mode of inter-macromolecular interaction under environmental conditions where these amino acids were sparse. The phenomenon of RNA-protein interaction is often considered central to early life and thought to predate LUCA and evolution of the full alphabet. Thus, the mechanism of how the early protein-RNA interaction could have been established in the absence of aromatic/basic residues has been debated. Besides a potential role of positively charged non-canonical amino acids (such as diaminobutyric acid) that could be present in the prebiotic environment, a hypothesis of negatively charged amino acids interacting with RNA via metallic cations (Mg^2**+**^ or Fe^2**+**^) has been raised (Vázquez-Salazar and Lazcano 2018)(Raggi et al. 2016). Depending on the abundance of positive and negative amino acids in the environment, both modes of interaction could have been present while experimental data to support such hypothetical modes of interaction are lacking. Importantly, a tRNA-binding peptide composed of only Gly, Ala, Asp and Val was recently selected using cDNA display but the structural mechanism of the binding has not been further characterized (Kumachi et al. 2016).

To shed more light on the possible mechanism of early protein-RNA interaction, we expressed the most enriched variants from the E and M pools of the CL11 protein domain and characterized them in both uncomplexed and complexed forms. Interestingly, both CL11-M and -E variants are less structurally ordered than the wild-type protein in their free form. This is probably caused by the lowered number of internal stabilizing interactions which in the case of the -E variant may mainly be due to a complete lack of aromatic amino acids (Longo et al. 2015; Makarov et al. 2021). Nevertheless, both CL11 variants bind to the cognate rRNA with an overall similar affinity *(K*_*D*_ in the order of 10^−9^ M), even though the different kinetics of the binding suggests additional folding of the proteins upon binding and/or a different mode of interaction. In the case of the CL11-E variant, the binding kinetics are significantly slower when compared with the wild-type CL11, suggesting that while the interaction is difficult to form, it is also more persistent.

However, the CL11-E/58rRNA complex stability is more sensitive to physicochemical conditions, such as temperature and pH. This feature was not recorded for the CL11-M/58rRNA complex which expressed an identical stability profile as the wild-type complex (Figure S4), suggesting that the binding mode is consistent when CL11 variants have all the positively charged and aromatic amino acid residues conserved. Most significantly, the CL11-E/58rRNA complex becomes unstable when salt is removed from the buffer composition, yet the absence of salt does not affect the secondary structure properties of the protein (Figure S3). This finding provides a hint that K^+^/Mg^2+^ ions may be important for the formation of the CL11-E/58rRNA complex while the overall structural shape of the complex is preserved with respect to the wild-type as suggested by the complex native ion mobility experiment. We further compared the CL11 and CL11-E RNA-binding properties using MD simulations. 30 direct H-bond interactions have been identified at the interface of wild-type CL11 and the 58rRNA, 12 of them mediated by Arg side chains. The latter interactions were obviously absent from the CL11-E binding interface. Further, water-mediated H-bonds via Lys side chains in CL11 were functionally compensated in CL11-E by water-mediated H-bonds via Ser side chains and Gly backbones. A unique feature of the CL11-E/58rRNA complex was the presence of K^**+**^/Mg^2**+**^ ion bridges found between acidic Glu side chains and the RNA phosphate backbone, which indicates the possibility of a new compensatory interaction mode between the protein and RNA (Figure 4). Although the MD simulation results are in line with the experimental observation of the CL11-E/58rRNA complex instability under no salt (no bridging cation) and low pH condition (protonated glutamic acid residues have a significantly reduced possibility to form the cation bridge interaction), the atomistic details of the binding interface should not be overinterpreted. Further investigations with MD as well as NMR, which are beyond the extent of this study, would shed more light on the studied interaction (Campagne et al. 2019). Previously, tens-of-microsecond-long or enhanced-sampling MD simulations proved useful in studying RNA-protein interfaces, including H-bonding, water and ion bridges (Krepl et al. 2017; Bochicchio et al. 2018). Additionally, free-energy simulation methods may help reveal the affinity contributions of individual protein residue mutations in RNA-protein complexes (Krepl et al. 2017; Bochicchio et al. 2018). But still, force-field parameters for RNA may be problematic in some cases and are under constant development via comparison with quantum mechanics (QM) calculations (Bochicchio et al. 2018; Pokorná et al. 2018).

To the best of our knowledge, this is the first experimental demonstration of cation mediated RNA-protein interaction, obtained via an *in vitro* evolution approach. Proteins lacking positively charged and aromatic residues may represent features of early proteins that could have existed during the pre-LUCA period. The structural and functional properties of early proteins are still debated. While previous studies have pointed out that acidic proteins can be soluble and sustain secondary structure information, it has generally been unclear how those proteins can form compact architectures and/or interact with the highly acidic nucleic acids (Doi et al. 2011; Tanaka et al. 2011; Longo et al. 2013; Shibue et al. 2018; Newton et al. 2019; Solis 2019). A recent study by the Tawfik group suggested that polyamines and divalent cations may have played an important role in promoting the folding of such early proteins (Despotović et al. 2020). Similarly, magnesium cations clearly play a special role in the ribosomal structure, a molecular fossil that reports on the earliest interactions between RNA and proteins (Petrov et al. 2011). In the ribosomal central and most conserved region, magnesium cation has even been observed to mediate RNA-protein (protein L2) interaction via water molecules (Petrov et al. 2012). More recently, RNA has been found to be generally stabilized by amino acid-chelated Mg^2^**+**, implying possibly a very fundamental role of metal cations in the coevolution of the RNA-protein world (Yamagami et al. 2018). Finally, our study suggests that such a mechanism of metal-ion assisted interaction could have enabled early RNA-protein interaction in the environment where positively charged amino acids were sparse. While further examples are needed, our case study suggests the possibility that positively charged amino acids were added later to the protein alphabet to fine-tuning of RNA-protein transient interactions but were not explicitly necessary either.

## Materials and Methods

### Electrophoretic mobility shift assay (EMSA)

Protein-RNA complexes were analysed following Xing *et al*. protocol (Xing and Draper 1995). Fluorescently labelled 58rRNA (f58rRNA) target was synthesized by Integrated DNA Technologies (Supplementing material). Purified proteins were incubated in 1:1 molar ratio with f58rRNA target (2 µM concentration of both) in buffer R (30 mM Tris, 20 mM MgCl_2_ and 175 mM KCl at pH 7.9) for 1 hour at room temperature. Gels were visualized by fluorescence in Typhoon FLA 9500.

### Surface Plasmon Resonance (SPR) Measurements

All SPR measurements were performed on a four-channel SPR sensor platform (PLASMON IV) developed at the Institute of Photonics and Electronics (IPE) of the Academy of Sciences of the Czech Republic, Prague. Gold SPR chips were functionalized following the Neburkova *et al* (Neburkova et al. 2018). protocol. Afterwards, 500 µL of 0.3 µM biotinylated target 58rRNA in buffer R + 0.1% Triton X-100 solution was loaded on the functionalized chip. Assay with immobilized neutravidin without 58rRNA target served as negative control to all kinetic experiments. CL11 (16 to 2 nM), CL11-M (30 to 3.75 nM) and CL11-E (580 to 285 nM) in buffer R + 0.1% Triton X-100 were injected (association phase) for several minutes, and then buffer R + 0.1% Triton X-100 alone was injected (dissociation phase). Obtained data were fitted by the logistic equation using TraceDrawer (Ridgeview Instruments AB), and K_on_, K_off_ and K_D_ values were calculated from Kinetic evaluation using OneToOne fitting model (Figure S4A).

### Temperature, pH and salts stability

Magnetic Dynabeads MyOne Streptavidin C1 (Invitrogen) and Magnetic Dynabeads M-280 Tosylactivated (Invitrogen) functionalized with Neutravidin (Invitrogen) were used to immobilize 1 μM of biotinylated 58rRNA according to the manufacturing recommendation. The functionalized beads were washed and then incubated for 1 hour with 1 μM solution of different protein variants in Buffer R + 0.05% Triton X100 at room temperature under gentle rotation. 20 μL (5 mg/mL) beads retaining the formed complex were washed and resuspended in buffer R + 0.05% Triton X-100 at the final concentration of 5 mg/mL and incubated for 10 minutes at different ranges of temperature (20-80ºC). In the case of pH stability, the beads were incubated 1 hour at room temperature at 3 different pH buffers: 50 mM Glycine, 175 mM KCl, 20 mM MgCl_2_, 0.05% Triton X-100 pH 3.5 or 10.5 and buffer R. As regarding the complex salts stability 5 mg/mL of functionalized beads were incubated 1 hour at room temperature in: buffer R, buffer R without KCl, buffer R without MgCl_2_, buffer R with 40 mM CaCl_2_ instead of MgCl_2_ and buffer R without KCl and MgCl_2_. Eluted samples were loaded on 16% SDS-PAGE gel and analyzed by Western blots using Monoclonal Anti-6X His tag antibody produced in mouse conjugated with HorseRadish Peroxidase, HRP (Sigma), detected with a Immobilon Forte Western HRP substrate (Merck) and visualize by Amersham Imager 600.

### Circular Dichroism

The ECD spectra were measured using the Jasco 1500 spectropolarimeter equipped with the Peltier holder PTC-517 in the far-UV (195-280 nm) spectral range. The ECD spectra were collected in different buffers composition (50 mM Glycine, 175 mM KCl, 20 mM MgCl_2_, pH 3.5 or 10.5, buffer R and buffer R without KCl and MgCl_2_) and in temperature range 5 °C to 95 °C with the step of 5 °C in 0.2 mm path length quartz cell (scanning speed of 10 nm/min, response time of 8 seconds, scanning step 0.5 nm) and with a samples concentration in buffer R of 0.2 mg/mL. After baseline correction, the final spectra were expressed as molar ellipticity (*q*) (deg.cm^2^.dmol^-1^) per residue. The numerical analysis of secondary structure was performed using the CDPro software package (Sreerama and Woody 2000)(Sreerama and Woody 2004).

### Molecular dynamics simulation

Starting from CL11/58rRNA complex (PBD: 1HC8, chains A,C) (Conn et al. 2002), the CL11-E/58rRNA complex was modelled by replacing the amino acids in question in PyMol, version 1.7.4 (The PyMOL Molecular Graphics System, Version 1.7.4, Schrödinger, LLC). Ions important for the structural stability of the RNA (Mg^2^**+** except residues 1165, 1166, two) Os^3+^ which were replaced by Mg^2^**+** for simplicity and one K^+^ were retained in the model. Apo proteins were modelled by deleting the RNA component. Water molecules were added extending 12 Å from the solute. Mg^2+^ and Cl^**-**^ ions were added to a final concentration of 20 mM and further K**+** and Cl^**-**^ ions were added to a final concentration of 175 mM. Mg^2+^ parameters of Allner *et al*. (Allnér et al. 2012) were used. The systems were stepwise minimized, heated to 300 K and at a pressure of 1 atm in NpT ensemble, production runs of molecular dynamics (MD) ensued for 2 μs. Frames were saved every 1 ns. The SHAKE algorithm was used to restrain all bond vibrations and hydrogen mass repartitioning to 3 Da allowed us to apply a time step of 4 fs. All the analyses were done in the CPPTRAJ programme (Shitov et al. 1984).

## Acknowledgments

We thank Dr. Milan Kožíšek for technical support with the SPR experiment, Dr. Radko Souček for the amino acid analysis, and Prof. Stephen Fried for his valuable comments on the manuscript. This work was supported by the Czech Science Foundation (GAČR) grant number 17-10438Y, the Human Frontier Science Program grant HFSP-RGY0074, European Commission H2020 project (EPIC-XS - grant agreement ID: 823839) and by the project “BIOCEV” CZ.1.05/1.1.00/02.0109. ML was supported by the project ‘Chemical Biology for Drugging Undruggable Targets’ (ChemBioDrug CZ.02.1.01/0.0/0.0/16_019/0000729) from the European Regional Development Fund (OP RDE), by the institutional project RVO 61388963 and by the Ministry of Education, Youth and Sports of the Czech Republic from the Large Infrastructures for Research, Experimental Development, and the Innovations project ‘IT4 Innovations National Supercomputing Center – LM2015070’. K.F. is supported by ELSI-First Logic Astrobiology Donation Program.

