## Supplementary material for "In vitro evolution reveals primordial RNA-protein interaction mediated by metal cations"

Klára Hlouchová

Supplementary Information Text

**Oligonucleotide sequences**

**>Forward E library**

5`-AAAAAGTGGCCTGTAATACGACTCACTATAGGGACACCAATAGAGAAAGAGGAGAAATACTAGATGGACTATAAAGATGACGACGATAAGACCGBAATTACCRGCACCCCGCCGGCGGCGGTGCTGCTGWYAGMCGCGGCGGGCATTGAAAGCGGCAGCGGCGAACCG-3`

**>Reverse E library**

5`-TTTTTCACCTGATCCGCTGCCATCTTCCACCACAATGCCTRHGCTTRYCGCGGTGCCTTCAATGVTTRHTRYCGCCGCTTCAATGCTCGCCGCTRKCAGATCCGGTRYTRHCAGTTCCGCAATTTCTKCCACTRHATCTABGSTAATGGTCGCCACTAYKKCTWCTHCCGGTTCGCCGCTGCCGCTTTCAATGCCCGCC-3`

**>Forward M library**

5`-AAAAAGTGGCCTGTAATACGACTCACTATAGGGACACCAATAGAGAAAGAGGAGAAATACTAGATGGACTATAAAGATGACGACGATAAGACCTTTATTACCAAAACCCCGCCGGCGGCGGTGCTGCTGAAAAAAGCGGCGGGCATTGAAAGCGGCAGCGGCGAA-3`

**>Reverse M library**

5`-TTTTTCACCTGATCCGCTGCCATCTTCCACCACAATGCCGVBGCTGCGCGCGGTGCCTTCAATTNYGCGTNYCGCCGCTTCAATGCTCGCCGCGVBCAGATCCGGTNYTTTCAGTTCCGCAATTTCGCGCACTTTATCGCGTTTAATGGTCGCCACTTTGVBGCGGVBCGGTTCGCCGCTGCCGCTTTCAATGCCCGCC-3`

**>RT-PCR Forward primer**

5`-AAAAAGTGGCCTGTAATACGACTCACTATAGGGACACCAATAGAGAAAGAGGAGAAATACTAGATGGACTATAAAGATGACG-3

**>RT-PCR Reverse primer**

5’-TTTTTCACCTGATCCGCTGCC-3’

**>Gateway Forward primer CL11-E**

5`-GGGGACAAGTTTGTACAAAAAAGCAGGCTTCAAAGAGGAGAAATACTAGATGCATCACCATCACCATCACACCGTTATCACCGGTACCCCG-3`

**>Gateway Forward primer CL11-M**

5`-GGGGACAAGTTTGTACAAAAAAGCAGGCTTCAAAGAGGAGAAATACTAGATGCATCACCATCACCATCACACATTCATCACCAAGACACCC-3`

**>Gateway Reverse primer CL11-E and CL11-M**

5`-GGGGACCACTTTGTACAAGAAAGCTGGGTCTTTTTCACCTGATCCGCTGCCTTA-3`

**>CL11 DNA sequence**

5`-ATGCATCACCATCACCATCACACCTTCATCACCAAAACCCCGCCGGCTGCTGTTCTGCTGAAAAAAGCTGCTGGTATCGAATCTGGTTCTGGTGAACCGAACCGTAACAAAGTTGCTACCATCAAACGTGACAAAGTTCGTGAAATCGCTGAACTGAAAATGCCGGACCTGAACGCTGCTTCTATCGAAGCTGCTATGCGTATGATCGAAGGTACCGCTCGTTCTATGGGTATCGTTGTTGAAGACTAA-3`

**>CL11-E DNA sequence**

5`-ATGCATCACCATCACCATCACACCGTTATCACCGGTACCCCGCCGCCGGCTGTTCTGCTGTCTGACGCTGCTTCTGTTGAATCTGGTTCTGGTGAACCGGAAGTTGAAGTTGTTGCTACCATCACCGTTGACCTGGTTGCTGAAATCGCTGAACTGGTTACCCCGGACCTGATCGCTGCTTCTATCGAAGCTGCTATCATCTCTATCGAAGGTACCGCTGCTTCTTCTGGTATCGTTGTTGAAGACTAA-3`

**>CL11-M DNA sequence**

5`-ATGCATCACCATCACCATCACACATTCATCACCAAGACACCCCCGGCGGCAGTTTTATTAAAAAAAGCTGCCGGCATTGAGTCAGGCAGCGGGGAACCTGGGCGTTTGAAAGTTGCTACGATTAAACGCGACAAGGTCCGTGAGATCGCTGAACTGAAACGCCCTGACTTGTCAGCGGCTTCGATCGAGGCAGCGATCCGTACAATCGAAGGGACCGCCCGCAGCGTAGGCATCGTTGTCGAAGATTAA-3`

**Puromycin complementary sequence**

5`-GGCAGCGGATCAGGTGAAAAA-3`

**Puromycin tag fragment**

5`-pCCCTTCACCTGATCCGCTGAAAAAAAAAAAAAAAAAA-(Spacer18)-(Spacer18) (Fluorescein-dT)-(Spacer18)-CC-(Puromycin)-3`

**Biotinylated 58rRNA target**

5`-biotin-(TEG)-GCCAGGAUGUAGGCUUAGAAGCAGCCAUCAUUUAAAGAAAGCGUAAUAGCUCACUGGU-3`

**Fluorescent 58rRNA target**

5`-fluorescein-(Spacer9)-GCCAGGAUGUAGGCUUAGAAGCAGCCAUCAUUUAAAGAAAGCGUAAUAGCUCACUGGU-3.

Supplementary methods

**mRNA display of CL11 combinatorial libraries and selection of RNA binders**. Each library M and E were prepared by Klenow extension of two overlapping single-stranded DNA oligonucleotides. ssDNA oligonucleotides were synthesized and PAGE purified by Integrated DNA Technologies. Annealing was performed by heating complementary oligonucleotide mixture (2 μM final concentration of each) in NEB1 buffer provided with 200 μM dNTPs to 90ºC for 2 minutes and cooling down to 25ºC for 10 minutes. Annealed product was filled in by Klenow extension to form double-strand DNA (dsDNA) as follows: 10 U of Klenow polymerase (NEB) was added to annealed oligonucleotides, incubated for 5 minutes at 25°C, 37°C for 1 hour and at 50°C for 15 minutes. Final dsDNA libraries were further column purified using the DNA Clean and Concentrator kit (Zymo Research). In the following transcription, 1μg of DNA library was used as a template for mRNA synthesis by T7-Flash kit (Lucigen). mRNA was purified by NucleoSpin® RNA Clean-up Kit (Macherey-Nagel) and used for ligation with puromycin tagged oligonucleotide fragment synthesized by Integrated DNA Technologies *via* T4 RNA ligase reaction (for details see Reyes *et al.* 2021(Reyes et al. 2021)). Ligated products were purified (final yield ~30%) and used for translation of protein-mRNA conjugated library, following the mRNA display protocol (Figure S1D)(Reyes et al. 2021). To minimize false positives, 300 to 400 ng mRNA-protein conjugates were mixed with pre-equilibrated magnetic Dynabeads MyOne Streptavidin C1 (Invitrogen) in buffer R (30 mM Tris, 20 mM MgCl_2_ and 175 mM KCl at pH 7.9) at 25°C with gentle mixing for 40 minutes, twice. The unbound products in the flow-through were collected and incubated for 1 hour at 25°C with 200 nM of the biotinylated 58rRNA target (Supplementing material) synthesized by Integrated DNA Technologies, Inc. The binding step was followed by biotinylated 58rRNA immobilization with bound protein library to 4.5 μL of pre-equilibrated streptavidin magnetic beads in buffer R and then washed 3-8 times with 500 μL of buffer R + 0.1% Triton X-100. Selected variants were eluted by heating the streptavidin affinity matrix for 2 minutes at 90°C in 5 μl of RNase free water. Total elution volume was used as a template for reverse transcription reaction (OneTaq® One-Step RT-PCR Kit, NEB) using previously described methods (Reyes et al. 2021). Reverse transcribed cDNA served as a template for the following selection rounds. mRNA display was carried out for 16 rounds for E library and 10 rounds for M library. Enriched DNA pools from selected rounds were submitted to next-generation sequencing via the Illumina MiSeq platform.

**Sequencing and data analysis.** Rounds 6, 9, 11, 14 and 16 of library E selection and rounds 1, 4 and 10 of library M were analyzed by high throughput sequencing on Illumina MiSeq. Prior to sequencing library preparation, quantification was carried out on Quantus™ Fluorometer (Promega). Total 100 ng of DNA sample was used as an input for the library preparation by NEBNext Ultra II DNA Library Prep kit (New England Biolabs) with AMPure XP purification beads (Beckman Coulter). Length of the prepared library was determined by Agilent 2100 Bioanalyzer (Agilent Technologies), quantified by Quantus Fluorometer (Promega). Sample was sequenced on MiSeq Illumina platform using the Miseq Reagent Kit v2 500-cycles (2x250) in a paired-end mode. Raw data was processed with Galaxy platform and sequence analysis of assembled and filtered paired reads was performed with MatLab scripts developed at Heinis lab (Rebollo et al. 2014; Afgan et al. 2018).

**Cloning, expression and purification of CL11, CL11-M and CL11-E.** The CL11, CL11-M and CL11-E genes were synthesized with N-terminal 6X Histidine tag by Invitrogen and amplified by PCR and cloned into expression vector pDEST14 (Invitrogen) *via* Gateway cloning system (Gateway Technology). The cloned genes were used to transform OverExpress C41(DE3) pLysS chemically competent *E. coli* (Invitrogen). Fresh transformants were suspended in 10 ml of sterilized ampicillin-supplemented (100 μg/ml) LB-medium and grown overnight at 37ºC and 220 rpm. 2 ml of the pre-culture were inoculated into 2 L of sterile LB-medium supplemented with ampicillin (100 μg/ml) and grown in a rotary incubator (Innova 4300, New Brunswick Scientific) at 37ºC and 220 rpm. Expression was induced after OD_600_ reached a value of ~0.9 by addition of IPTG to a final concentration of 1 mM. Cells were cultivated for further 3.5 hours at 25ºC and 200 rpm and harvested by centrifugation (6000 rpm, 10 minutes, 4ºC). Pellets were resuspended in B-PER Bacterial Protein Extraction Reagent (Thermo Fisher Scientific) at 2.5 mL/g wet cells pellet with 1.5 U/mL of RNase A and DNAse I (Jena Bioscience) and protease inhibitor (Sigma). Cell suspension was incubated at room temperature for 20 minutes on a vascular device at 60 rpm. Cells were centrifuged at 18,000 rpm for 20 minutes. The supernatants were loaded on 1 mL of Talon Metal Affinity Resin (Clontech Laboratories, Inc.) equilibrated with Talon buffer A (300 mM NaCl, 50 mM NaH_2_PO_4_, 50 mM of Glutamic acid salt at pH 7.2) and incubated under agitation for 30 minutes at room temperature. Column was washed with Talon buffer B (300 mM NaCl, 50 mM NaH_2_PO_4_, 50 mM of glutamic acid salt, 5 mM imidazole at pH 7.2). Samples were eluted by 2 mL of Talon buffer C (300 mM NaCl, 50 mM NaH_2_PO_4_, 50 mM of glutamic acid salt, 250 mM imidazole at pH 7.2). The CL11 and CL11-M fractions were dialyzed and loaded on HiTrap CaptoS 5 mL cation exchange column (GE Healthcare) equilibrated with buffer A (10 mM NaH_2_PO_4_ and 2 mM KCl at pH 7). The column was washed with five column volumes of buffer A. Bound proteins were eluted with 0-50% gradient of buffer B (10 mM NaH_2_PO_4_, 2 M NaCl and 2 mM KCl at pH 7). Fractions containing proteins of interest were collected, concentrated and loaded on Superdex 75 10/300 gel filtration column (GE Healthcare) equilibrated in a buffer containing 30 mM Tris, 20 mM MgCl_2_ and 175 mM KCl at pH 7.9. CL11 and CL11-M fractions were collected, concentrated, dialyzed in buffer R and stored at -80ºC. CL11-E eluate from Talon affinity purification was collected and incubated in Talon buffer A with 7 M urea for 16 hours at room temperature. The sample was loaded on 0.5 mL of Talon Metal Affinity Resin equilibrated with Talon buffer A + 7M urea and incubated for 2 hours at room temperature. Column was washed with Talon buffer B + 7M urea. Sample was eluted by 0.650 mL of Talon buffer C + 7M urea. The eluted sample was loaded on HiLoad 16/600 Superdex 75 gel filtration column (GE Healthcare) equilibrated in a buffer R. Fractions containing CL11-E were collected, concentrated, dialyzed in buffer R and stored at -80 ºC. Protein concentrations were determined by amino acid analysis using a Biochrom 30+Series Amino Acid Analyser (Biochrom, UK).

**Native ion mobility - mass spectrometry (IMS-MS) measurements**. Native IMS-MS experiments were conducted on Waters Synapt G2Si instrument equipped with traveling wave IMS (TWIMS). The complex was assembled mixing 27 µL of 30 µM CL11 or CL11-E sample with 3 µL 300 µM RNA in buffer R. Immediately prior MS analysis, samples were buffer exchanged to ammonium acetate (150mM, pH 6.9) using Zeba™ Micro Spin Desalting Columns (7K MWCO, 75 µL, Thermo Fisher Scientific). Desalted samples (25µM protein-58rRNA complexes) were loaded into quartz nESI emitters pulled on P‑2000 laser puller (Sutter instruments). Measurement was performed in negative ion mode for mass range 500-5000 Da with IMS wave velocity 750 m/s and height 30 V. For sample ionization ESI voltage (0.5-1 kV) was applied to platinum wire placed into nESI emitter containing sample solution. Obtained data were analyzed using MassLynx 4.1 (Waters) and OriginPro2015 (OriginLab). Collision cross sections (CCS) evaluation was performed using logarithmic calibration on native proteins (Cyt C, UBQ and bLG)(Bush et al. 2010; Allen et al. 2016)

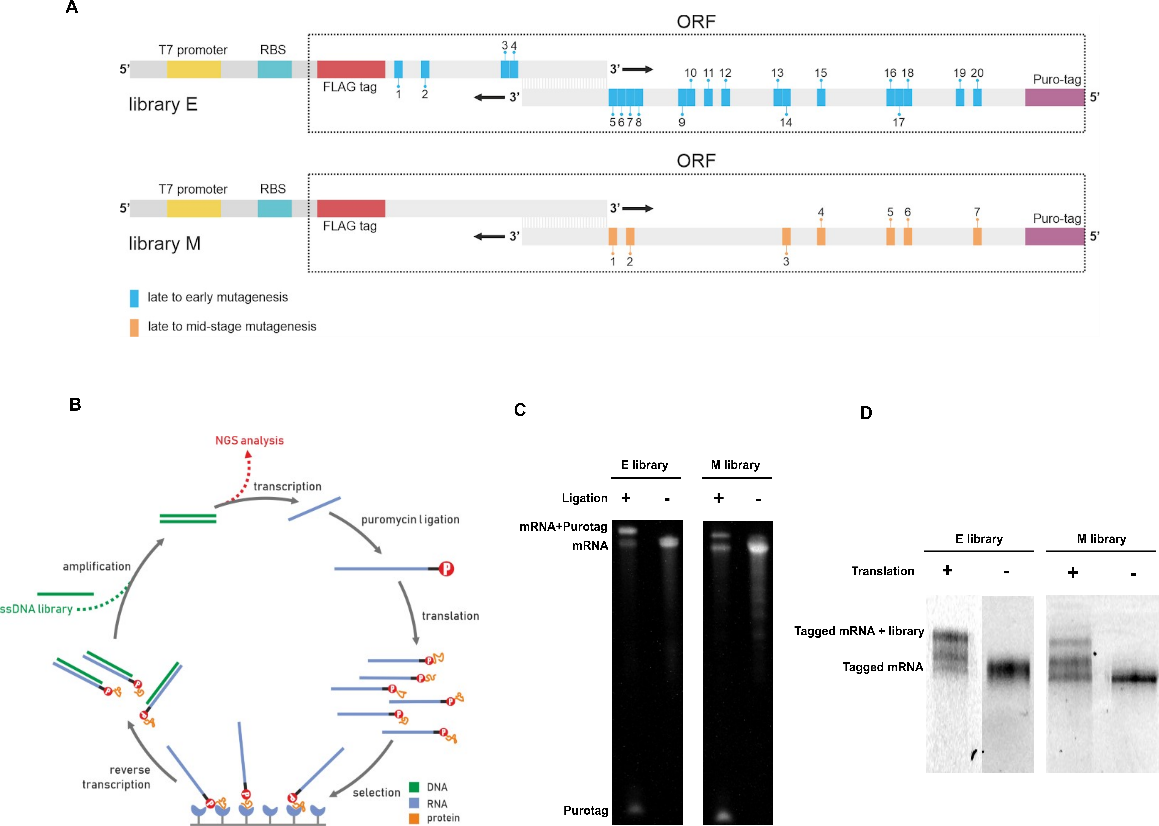

Fig. S1. mRNA display and preparation of the library E and M. (A) Design of the ssDNA libraries expression cassette (RBS - ribosomal binding site, ORF - open reading frame). (B) Schematic representation of the mRNA display and selection pipeline. The methodology starts with the annealing and the amplification of the designed ssDNA library. The DNA library is in vitro transcribed to mRNA library and undergoes ligation with the puromycin-DNA tag. Tagged product is translated and the resulting sample is mRNA-protein conjugate used for the selection process toward 58rRNA target. The selected variant conjugates are reverse transcribed and sequenced. (C)  Images of mRNA ligation with Purotag fragment for M and E library resolved in 8 % polyacrylamide TBE with 8 M Urea gels with GelRed staining. (D) Images of the translated tagged mRNA resolved in 8 % polyacrylamide TBE with 8 M Urea gels with fluorescent visualization.

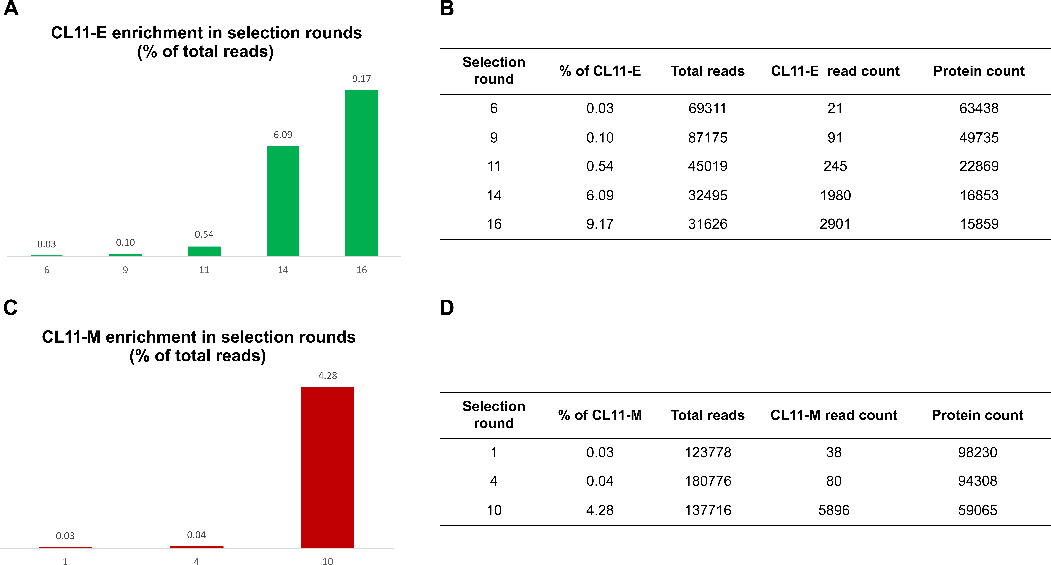

Fig. S2. Analysis of the mRNA display selection sequencing data (A) CL11-E enrichments from round 6th to 16th round selection. (B) Total E library enrichments from round 6th to 16th round selection. (C) CL11-M enrichments from round 1st to 10th round selection. (D) Total M library enrichments from round 1st to 10th round selection.

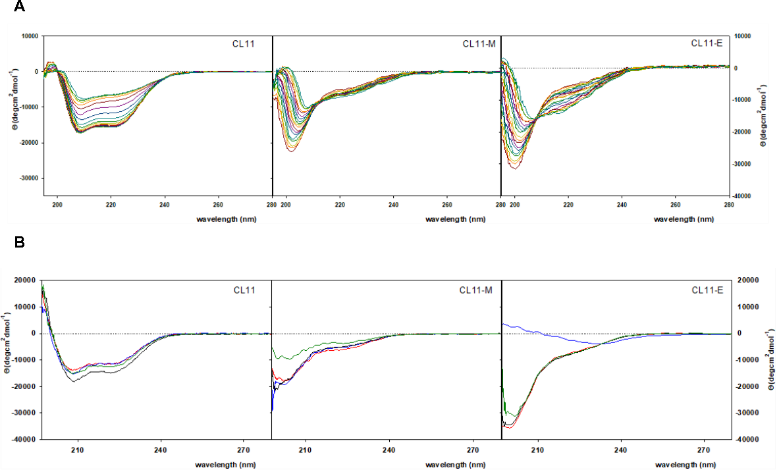

**Fig. S3.** **ECD spectra of protein CL11, CL11-E and CL11-M.** (A) ECD spectra of protein CL11, CL11-E and CL11-M in temperature interval 5-95°C. (B) The ECD spectra of CL11, CL11-E and CL11-M in buffer pH 7.9 (black), buffer pH 3.5 (blue), buffer pH 10.5 (green) and buffer pH 7.9 without Mg^2^**^+^** and K**^+^** (red).

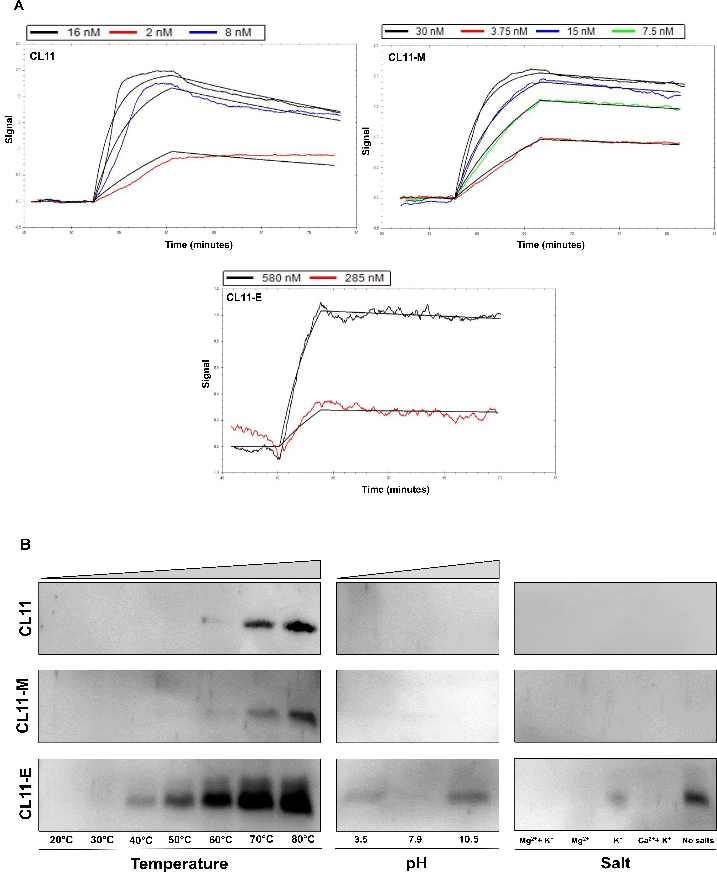

**Fig. S4. Complex characterization of the CL11, CL11-M and CL11-E.** (A) The biotinylated target RNA was immobilized on SPR chips through a neutravidin−biotin interaction. For each protein different concentration were injected (association phase): CL11 (from 16 nM to 2 nM), CL11-M (from 30 nM to 3.75 nM) and CL11-E (from 570 nM to 285 nM) and then Buffer R with 0.1% Triton X100 was injected (dissociation phase). (B) Western Blot of the protein-RNA complex stability at different temperatures, pHs and salts presence.

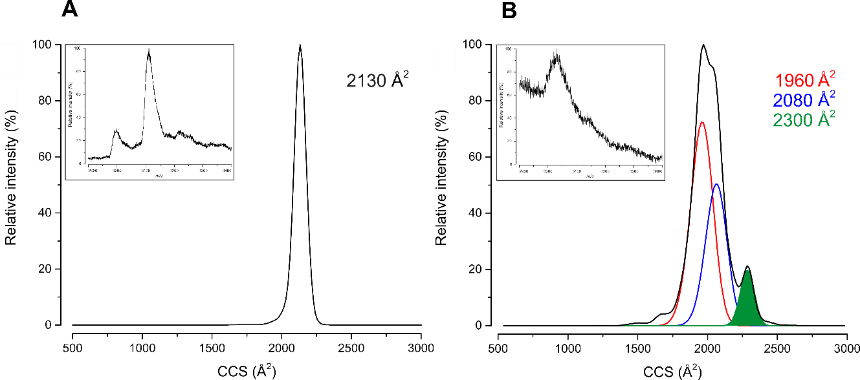

**Fig. S5.** Ion mobility trace for CL11-58rRNA complex (A) and CL11-E-58rRNA complex (B). The insets represent 9- charge states of protein-RNA complexes which were selected to derive IMS trace.

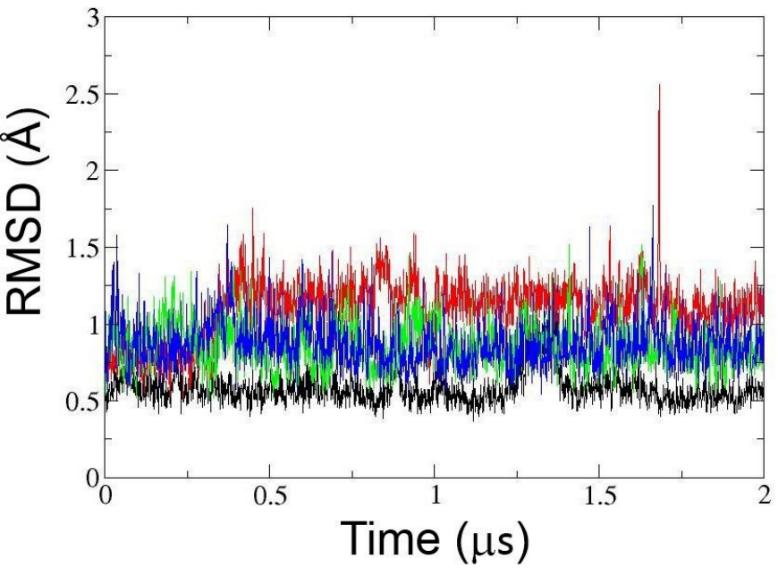

**Fig. S6.** RMSD of protein secondary structure backbone atoms during 2 µs MD of CL11-58rRNA (black), CL11-E-58rRNA (red), apo CL11 (green) and apo CL11-E (blue).

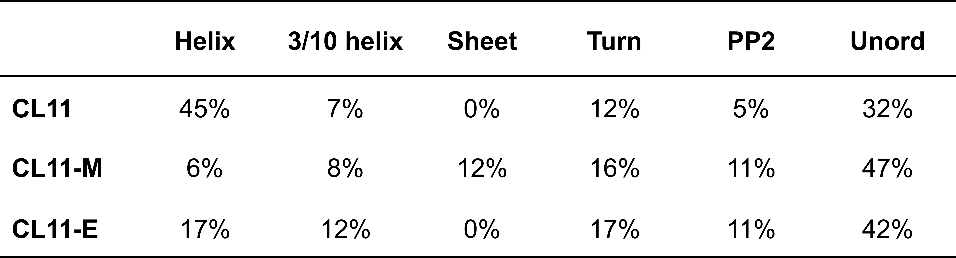

**Tab. S1. Analysis of the secondary structure content of CL11, CL11-M and CL11-E** revealed by numerical analysis by ECD spectra performed using the CDPro software package (Sreerama and Woody 2000; Sreerama and Woody 2004).

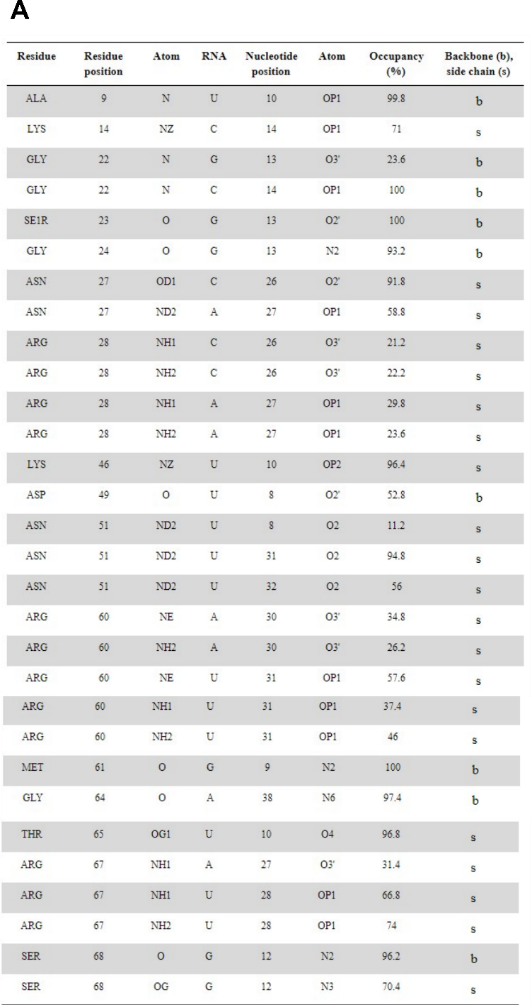

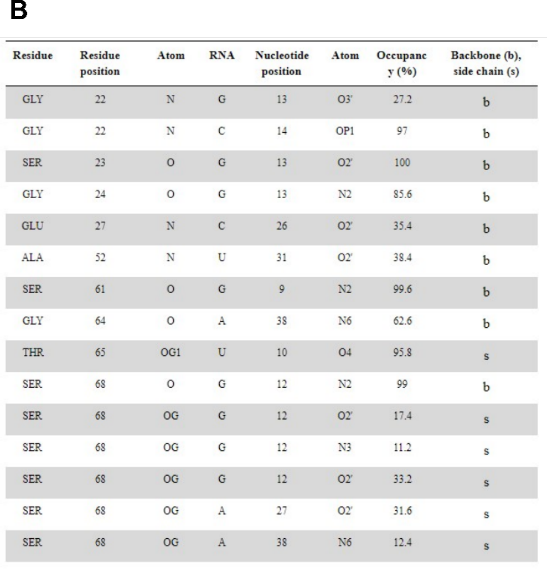

**Tab. S2**. Direct protein-RNA hydrogen bonding. (A) CL11/58rRNA, (B) CL11-E/58rRNA
